# *Leishmania* exploits the macrophage endoplasmic reticulum-shaping protein CLIMP-63 to modulate mitochondrial biogenesis and bioenergetics

**DOI:** 10.64898/2026.03.19.712868

**Authors:** Charles-Antoine Boyer, Hamlet Acevedo Ospina, Albert Descoteaux

## Abstract

In *Leishmania*-infected macrophages, interactions between parasitophorous vacuoles and the host cell endoplasmic reticulum are central to parasite replication. Whether the complex architecture of this organelle impacts these interactions remains however unknown. Here, we identify the macrophage endoplasmic reticulum-shaping protein CLIMP-63 as a host factor promoting *Leishmania* infection. Following parasite internalization, we observed a redistribution of CLIMP-63 to the vicinity of parasitophorous vacuoles in a process dependent on the virulence glycolipid lipophosphoglycan, and its dissociation from the mitochondrial network. Mechanistically, we found that CLIMP-63 is required for *Leishmania*-induced mitochondrial DNA replication, cristae biogenesis, and enhanced mitochondrial respiration, which are critical for the ability of the parasite to colonize macrophages. Collectively, our findings uncover a role for CLIMP-63 in supporting macrophage colonization by *Leishmania* and illustrate how this intracellular pathogen exploits the host cell endoplasmic reticulum to reprogram mitochondrial functions and bioenergetics.

**AUTHOR SUMMARY:** The protozoan parasite *Leishmania* replicates in macrophages within parasitophorous vacuoles. These vacuoles establish interactions with various organelles and modulate their properties and functionality to adapt the host cell to their own needs. In this regard, *Leishmania* rewires host cell mitochondria to produce key metabolites and to modify their bioenergetic profile. We obtained evidence that *Leishmania* exploits the role of the macrophage endoplasmic reticulum architectural protein CLIMP-63 to stimulate mitochondrial biogenesis and respiration. The fact that CLIMP-63 plays a central role in the ability of *Leishmania* to colonize macrophage illustrates how this parasite manipulates the host cell endoplasmic reticulum to modulatet the function and properties of other organelles.

## INTRODUCTION

During their life cycle, protozoan parasites of the genus *Leishmania* alternate between the midgut of their insect vector, where they proliferate as promastigotes, and mammalian macrophages, where they replicate as amastigotes within parasitophorous vacuoles (PVs). These specialized compartments display a dynamic and hybrid composition, reflecting their interactions with various host cell organelles. Interactions between *Leishmania*-harboring PVs (LPVs) and the macrophage endoplasmic reticulum (mER), which involve both vesicular trafficking and membrane contact sites, are crucial for LPV expansion and maintenance, and to satisfy the complex nutritional requirements and auxotrophies of the parasites (1–8). These interactions also enable the transfer of virulence factors from LPVs to the mER, including the abundant surface glycolipid lipophosphoglycan (LPG) and the zinc-dependent metalloprotease GP63 (8–10). Although these two glycoconjugates play a major role in the interaction of *Leishmania* with macrophages, the significance of their presence in the mER has received little attention. Whereas GP63 was shown to cleave the ER isoform of syntaxin-5 (11), the impact of LPG accumulating into the mER remains unknown.

The ER is the largest cellular organelle and is made of a continuous dynamic membrane system that constitutes half of the membranous content of the cell. This organelle is organized into three distinct domains: the nuclear envelope, smooth peripheral tubules, and ribosome-studded rough ER sheets (12). Key cellular processes such as lipid synthesis and carbohydrate metabolism take place in the smooth peripheral ER, whereas rough ER sheets are the site of protein synthesis (13). The complex architecture and topology of the ER domains are maintained by ER-shaping proteins including reticulon (RTN), deleted in polyposis 1 (DP1), atlastin, and the cytoskeleton-linking membrane protein 63 (CLIMP-63) (13–16). RTN and DP1 interact to shape the tubular ER by stabilizing the high curvature of the tubules (17, 18), whereas atlastin maintains the ER network by regulating homotypic fusion between tubules (19). CLIMP-63, an abundant ER type II transmembrane protein, is essential for ER sheet formation and structure. It forms coiled-coil homodimers that span the ER lumen where it acts as an ER sheet spacer and anchors the ER to microtubules on the cytosolic side (20, 21). RTN and DP1 are also enriched at the curved edges of ER sheets whereas CLIMP-63 is also expressed in peripheral tubules (18, 22). The relative abundance of ER sheets and tubules is cell type- and context-dependent and is regulated by the expression levels of these ER-shaping proteins (23). A growing body of evidence indicates that host cell ER-shaping proteins play a role in host-pathogen interactions, where their recruitment to pathogen-containing vacuoles may modulate intracellular replication. Hence, atlastin and RTN4 were shown to localize to *Legionella-*containing vacuoles and to promote intracellular bacterial growth (24, 25). A more recent example is the reduced proliferation of *Salmonella* observed in RTN4-depleted cells and in CLIMP-63-overexpressing cells, suggesting that optimal *Salmonella* replication is influenced by the relative abundance of ER sheets and tubules (26).

In addition to its structural role, CLIMP-63 functions as a regulator of ER-mitochondria contact sites (ERMCS) by interacting with the mitochondrial porin VDAC2 (27). Through these interactions, CLIMP-63 modulates mitochondrial DNA synthesis, mitochondrial and cristae morphology, as well as mitochondrial respiration (27–31), consistent with a functional link between ER architecture and mitochondrial function. This CLIMP-63 function is highly relevant in the context *L. donovani* infection, as stimulation of macrophage mitochondrial biogenesis and oxidative phosphorylation is critical for the ability of the parasite to colonize host cells (32–35).

Here, we investigated whether the architecture of the mER influences the ability of *Leishmania* to colonize macrophages. We identify CLIMP-63 as a host factor that contributes to parasite replication and show that *Leishmania* exploits this ER-shaping protein to reprogram host cell mitochondrial biogenesis and bioenergetics.

## RESULTS

### CLIMP-63 redistributes to *Leishmania*-harboring parasitophorous vacuoles

Interactions between *Leishmania*-harboring PVs (LPVs) and the mER are essential for *Leishmania* replication and LPV expansion (1, 3, 6, 8). However, the role of the mER architecture in influencing these interactions remains poorly understood. We therefore sought to examine the contribution of the mER-shaping proteins CLIMP-63 and RTN4 to the ability of *Leishmania* to colonize macrophages. We infected bone marrow-derived macrophages (BMM) with metacyclic promastigotes of either *L. donovani*, which replicates in tight-fitting individual PVs, or *L. amazonensis*, which develops in large communal PVs. At different time points post-phagocytosis, we assessed the distribution of CLIMP-63 and RTN4 by confocal immunofluorescence microscopy. As shown in Fig 1A and B, CLIMP-63 displayed a punctate distribution in uninfected BMM. At 2 h, 6 h, and 24 h post-phagocytosis, CLIMP-63 redistributed in close proximity to the PVs in BMM infected with either *L. donovani* or *L. amazonensis* (Fig 1A, B). We also observed that 24 h and 48 h post-phagocytosis with *L. amazonensis*, CLIMP63 progressively accumulated to the membrane of large communal PVs (Fig 1B). To rule out the possibility that the staining signal for CLIMP-63 in close proximity to parasites was artefactual or due to cross-reactivity with *Leishmania* molecule(s), we used siRNA to deplete CLIMP-63 in BMM prior to infection (Fig 1A and B). Absence of signal for CLIMP-63 confirmed the efficacy of the siRNA-mediated depletion of CLIMP-63 as well as the specificity of the anti-CLIMP-63 staining. Infection with either *L. donovani* or *L. amazonensis* had no significant effect on CLIMP-63 levels, indicating that *Leishmania* only altered its sub-cellular distribution (Fig 1C). In contrast to CLIMP-63, the punctate distribution of RTN4 was not modified following infection (S1 Fig A and B), suggesting a selective impact of *Leishmania* on the distribution of mER-shaping proteins. Insertion of the *Leishmania* virulence glycolipid LPG into host cell membranes alters membrane protein distribution (36–38). To determine whether LPG mediates the redistribution of CLIMP-63, we infected BMM with metacyclic promastigotes of the *L. donovani* LPG-defective Δ*lpg1* mutant and of its complemented counterpart Δ*lpg1*+*LPG1*. At 6 h post-phagocytosis, CLIMP-63 localized to LPVs in BMM infected with Δ*lpg1*+*LPG1*, whereas the Δ*lpg1* mutant failed to induce CLIMP-63 redistribution (Fig 2A), identifying LPG as the *Leishmania* factor responsible for this effect. Because CLIMP-63 associates with ERMCS (27), we examined its colocalization with the mitochondrial protein Tom20. Strikingly, the colocalization between CLIMP-63 and Tom20 observed in uninfected BMM was significantly reduced in *L. donovani*-infected BMMs, indicating that redistribution of CLIMP-63 coincides with its dissociation from the mitochondrial network (Fig 2B). Collectively, these results demonstrate that *Leishmania* induces an LPG-dependent redistribution of CLIMP-63 and its dissociation from the mitochondrial network, and suggest a re-organization of the mER architecture.

**Fig 1.**
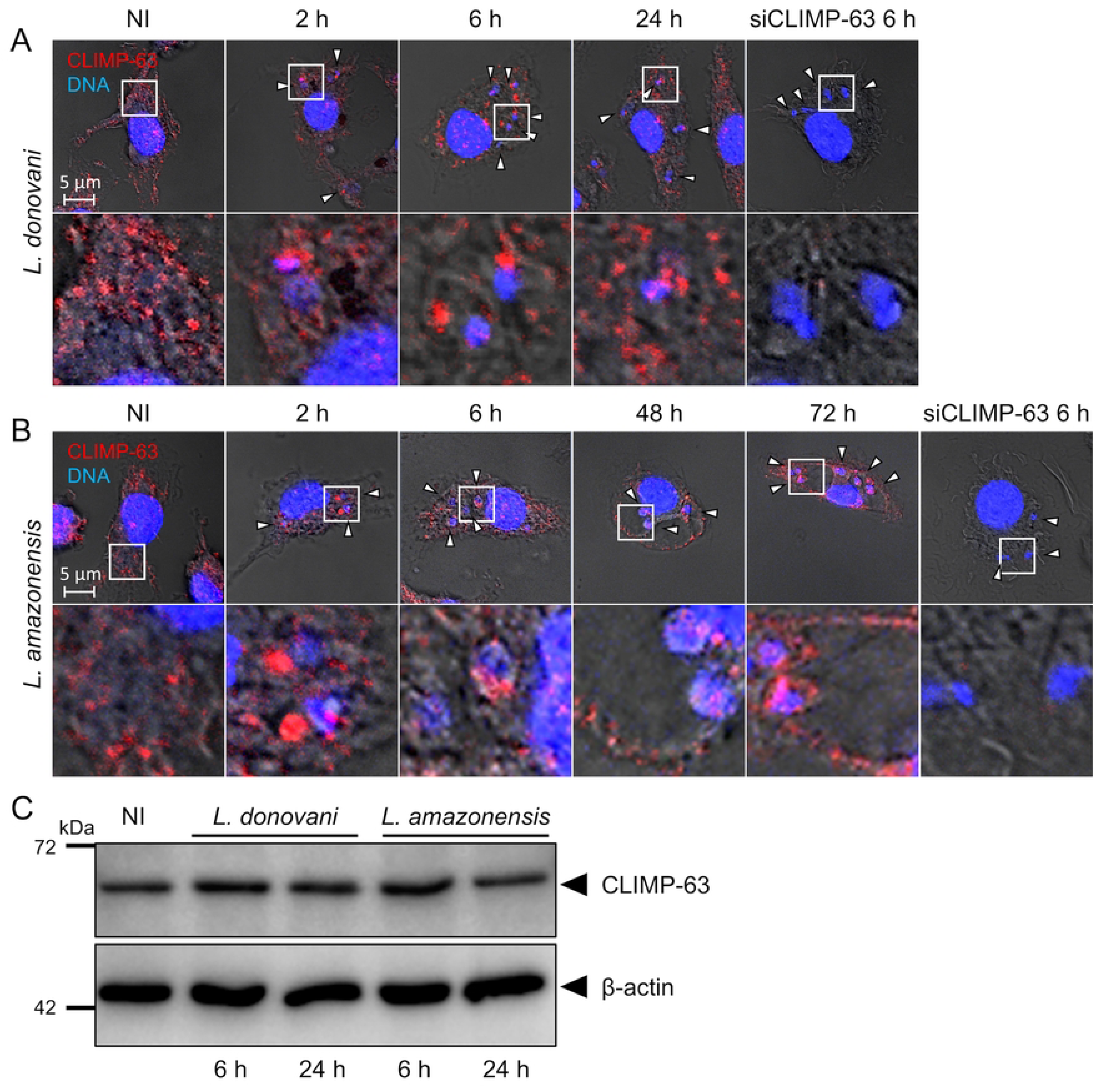
*Leishmania* induces the redistribution of CLIMP-63. BMM were infected or not (NI) with either (A) *L. donovani* or (B) *L. amazonensis* metacyclic promastigotes and at the indicated times points, the distribution of CLIMP-63 (red) was assessed by confocal immunofluorescence microscopy. DNA is in blue. White arrowheads denote internalized parasites. 5X-enlarged insets of representative localization of CLIMP-63 are shown. Controls for the specificity of the anti-CLIMP-63 antibody were performed using BMM pretreated with siRNA to CLIMP-63 and infected for 6 h with either *L. donovani* or *L. amazonensis* metacyclic promastigotes. Results are representative of at least three independent experiments. (C) Lysates of uninfected BMM (NI) and of BMM infected with either *L. donovani* or *L. amazonensis* metacyclic promastigotes were prepared at the indicated time points in lysis buffer containing 10 mM 1,10-phenanthroline. Levels and integrity of CLIMP-63 were assessed by Western blot analysis. β-actin was used a loading control. Immunoblots shown are representative of at least three independent experiments.

**Fig 2.**
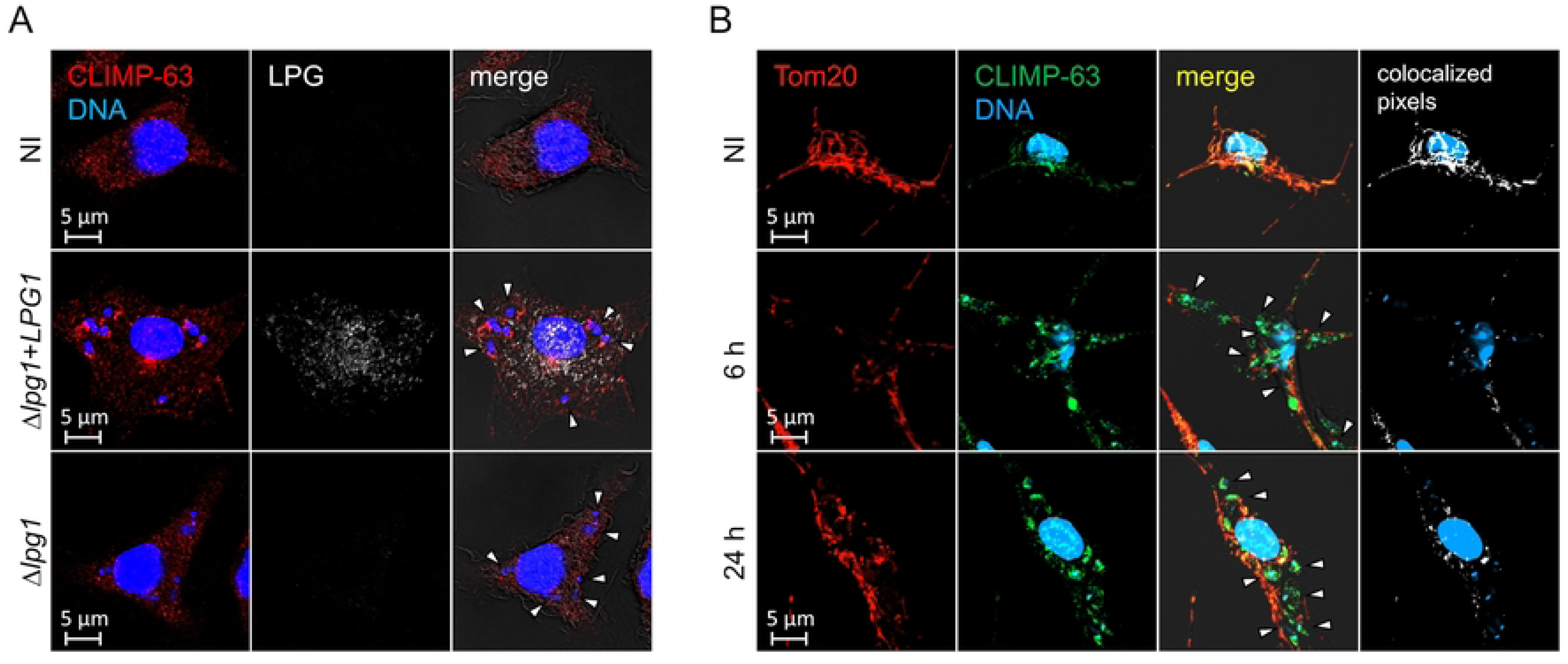
Redistribution of CLIMP-63 is LPG-dependent and coincides with its dissociation from the mitochondrial network. (A) BMM were infected or not (NI) with either *Δlpg1* or *Δlpg1+LPG1 L. donovani* metacyclic promastigotes and at the indicated times points the distribution of CLIMP-63 (red) was assessed by confocal immunofluorescence microscopy. DNA is in blue. White arrowheads denote internalized parasites. (B) BMM were infected with *L. donovani* metacyclic promastigotes and at the indicated times points the distribution and colocalization of CLIMP-63 (green) and Tom20 (red) was assessed by confocal immunofluorescence microscopy. Merge images and colocalized pixels (white) are shown. DNA is in blue. White arrowheads denote internalized parasites. Results are representative of at least three independent experiments.

### CLIMP-63 contributes to the ability of *Leishmania* to establish infection

The *Leishmania*-induced redistribution of CLIMP-63 and its dissociation from the mitochondrial network led us to examine its impact on parasite replication. Although *Leishmania* did not alter the distribution of RTN4 (S1 Fig), we also investigated its potential role in *Leishmania* replication. We infected BMM (treated with either control siRNA, siRNA to CLIMP-63, or siRNA to RTN4) with *L. donovani* or *L. amazonensis* metacyclic promastigotes and we assessed parasite burden up to 72 h post-infection. As depicted in Fig3A-B, depletion of CLIMP-63 did not affect the internalization of *L. donovani* and *L. amazonensis*. In *L. donovani*-infected BMM, parasite burden was reduced by 30% in CLIMP-63-depleted cells compared to controls during the first 24 h of infection, suggesting that CLIMP-63 contributes to the adaptation of *L. donovani* to the intracellular environment and to establish infection. Beyond 24 h, *L. donovani* replication was significantly impaired in CLIMP-63-depleted BMM compared to control BMM (Fig3A). In *L. amazonensis*-infected BMM, parasite burden was also reduced by 30% at 24 h post-phagocytosis in CLIMP-63-depleted BMM but in contrast to *L. donovani*, *L. amazonensis* replicated at a similar rate in control and CLIMP-63-depleted BMM (Fig3B). However, expansion of *L. amazonensis*-harboring communal PVs was significantly impaired in CLIMP-63-depleted BMM (Fig3C). To further assess the requirement for CLIMP-63 for the replication of *L. donovani*, we measured BrdU incorporation by intracellular parasites (8, 39). BMM (untreated or treated with siRNA to CLIMP-63 or with control siRNA) were infected with *L. donovani* metacyclic promastigotes and were incubated in the presence of BrdU for up to 72 h post-infection. As shown in S2 Fig, replication of *L. donovani* amastigotes, determined by the percentage of BrdU-positive parasites, was reduced by 50% at 48 h and 72 h post-infection in CLIMP-63-depleted BMM compared to controls, consistent with the impaired parasite replication observed in Fig 3A. In contrast to CLIMP-63, RTN4 played no significant role in *Leishmania* replication, nor in communal PVs expansion (Fig 3D-F). Collectively, these results demonstrate that CLIMP-63 is required for the replication of *L. donovani* and for the expansion of *L. amazonensis*-harboring communal PVs.

**Fig 3.**
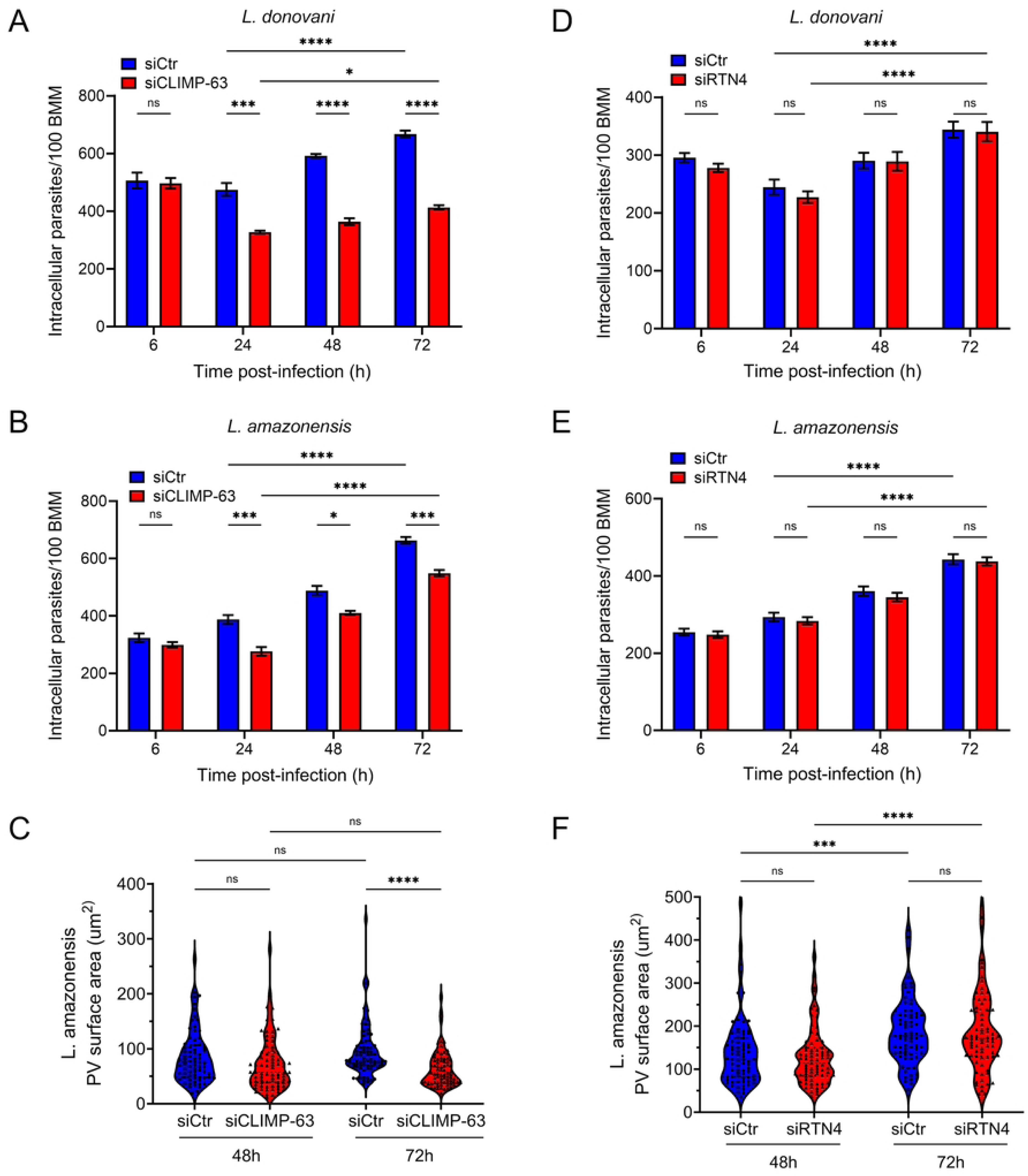
CLIMP-63 is required for the establishment of infection in BMM. BMM (treated with siRNA to either CLIMP-63 or RTN4 or treated with control siRNA, Ctr) were infected with *L. donovani* or *L. amazonensis* metacyclic promastigotes and at various time points post-phagocytosis parasite replication and PV size were assessed. (A) Quantification of *L. donovani* parasite burden in CLIMP-63 depleted BMM at 6, 24, 48, and 72 h post-infection. (B) Quantification of *L. amazonensis* parasite burden in CLIMP-63 depleted BMM at 6, 24, 48, and 72 h post-infection. (C) Quantification of PV size in CLIMP-63-depleted BMM infected with *L. amazonensis* at 48 and 72 h post-phagocytosis. (D) Quantification of *L. donovani* parasite burden in RTN4-depleted BMM at 6, 24, 48, and 72 h post-infection. (E) Quantification of *L. amazonensis* parasite burden in RTN4-depleted BMM at 6, 24, 48, and 72 h post-infection. (F) Quantification of PV size in RTN4-depleted BMM infected with *L. amazonensis* at 48 and 72 h post-phagocytosis. Data in (A, B, D, E) are presented as the means ± SEM of values of one representative experiment of three independent experiments. Data in (C, F) are presented as a violin plot with means ± standard deviations (SD) of values from three independent experiments for a total of 450 PVs. Statistics were calculated using one-way analyses of variance (ANOVA) with Sidak’s multiple comparison test with * P ≤ 0.05, *** P ≤ 0.001 and **** P ≤ 0.0001 significance. Blots showing the efficacy the siRNA-mediated CLIMP-63 and RTN4 knockdowns are shown in S3 Fig and S4 Fig, respectively.

### CLIMP-63 is required for *Leishmania*-induced modulation of mitochondrial biogenesis and metabolism

CLIMP-63 was reported to regulate mitochondrial DNA synthesis, mitochondria morphology, ER-mitochondria contact sites, as well as mitochondrial respiration (27–31). Given the importance for *Leishmania* to stimulate mitochondrial biogenesis and respiratory functions in host macrophages (32–34), we examined whether depletion of CLIMP-63 affects mitochondrial functionality in *L. donovani*-infected BMM. As illustrated in Fig 4A, the mitochondrial-to-nuclear (mt/n) DNA ratio was similar in uninfected control and CLIMP-63-depleted BMM. In contrast, at 24 h post-phagocytosis, *L. donovani*-induced mitochondrial DNA synthesis was abrogated in CLIMP-63-depleted BMM. AICAR, which stimulates mitochondrial biogenesis, was used as a positive control. Transmission electron microscopy analysis revealed no significant differences in the number of mitochondria in either uninfected or infected control and CLIMP-63-depleted BMM (Fig 4B, E). In contrast, the *L. donovani*-induced increase in the numbers of cristae per mitochondria and per cell observed at 24 h post-phagocytosis was abrogated in the absence of CLIMP-63 (Fig 4C-E). We next assessed the impact of CLIMP-63 on mitochondrial respiration using a Seahorse analyzer (Fig 5A). Consistent with previous reports (32, 34), *L. donovani* stimulated an increase in basal oxygen consumption rate, maximal respiration, non-mitochondrial oxygen consumption, as well as mitochondrial ATP production (Fig 5B-E). Importantly, the increase in mitochondrial respiration was impaired in CLIMP63-depleted BMM at 24h post infection, correlating with the reduced mitochodrial DNA synthesis, mitochondrial cristae and the reduced parasite burden observed in Figs 3 and 4. While CLIMP63 was required for the increase of the bioenergetic profile in *L. donovani*-infected BMM, proton leak remained unaffected, indicating that mitochondrial integrity is not affected (Fig 5F). Collectively, these results demonstrate that *L. donovani* requires CLIMP-63 to modulate host cell mitochondrial biogenesis and metabolism, which may contribute to the establishment of *L. donovani* infection within macrophages.

**Fig 4.**
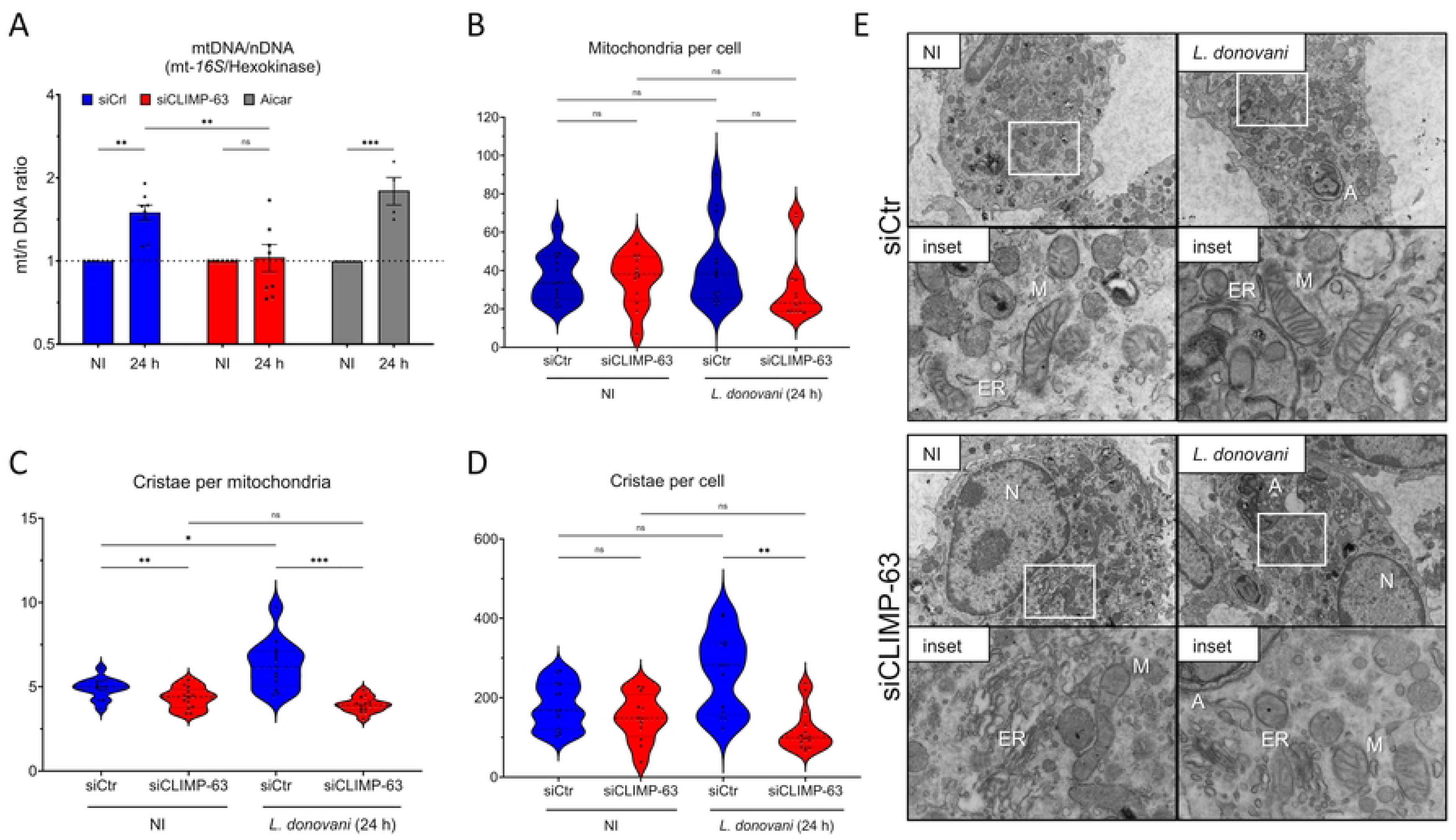
CLIMP63 is required for *L. donovani*-induced host cell mitochondrial biogenesis. (A) BMM (treated with siRNA to CLIMP-63 or treated with control siRNA, Ctr) were infected or not (NI) with *L. donovani* metacyclic promastigotes and at 24 h post-infection mitochondrial-over-nuclear (mt/n) DNA ratio (MT-16S/hexokinase) was determined. BMM were incubated for 24 h with 0.1 mM AICAR as a positive control. (B-E) BMM (treated with siRNA to CLIMP-63 or treated with control siRNA, Ctr) were infected or not (NI) with *L. donovani* metacyclic promastigotes and at 24 h post-infection cells were processed for electron microscopy (E). (B) The number of mitochondria per cell, (C) the number of cristea per mitochondrion and (D) the number of cristea per cell were determined from (E). The data are presented as mean values +/-SEM from three independent experiments. Statistics were calculated using one-way analyses of variance (ANOVA) with Sidak’s multiple comparison test for (A) and Tukey’s multiple comparison test for (B, C, D) with * P ≤ 0.05, ** P ≤ 0.01 and *** P ≤ 0.001 significance. Blots showing the efficacy the siRNA-mediated CLIMP-63 knockdowns are shown in S3 Fig.

**Fig 5.**
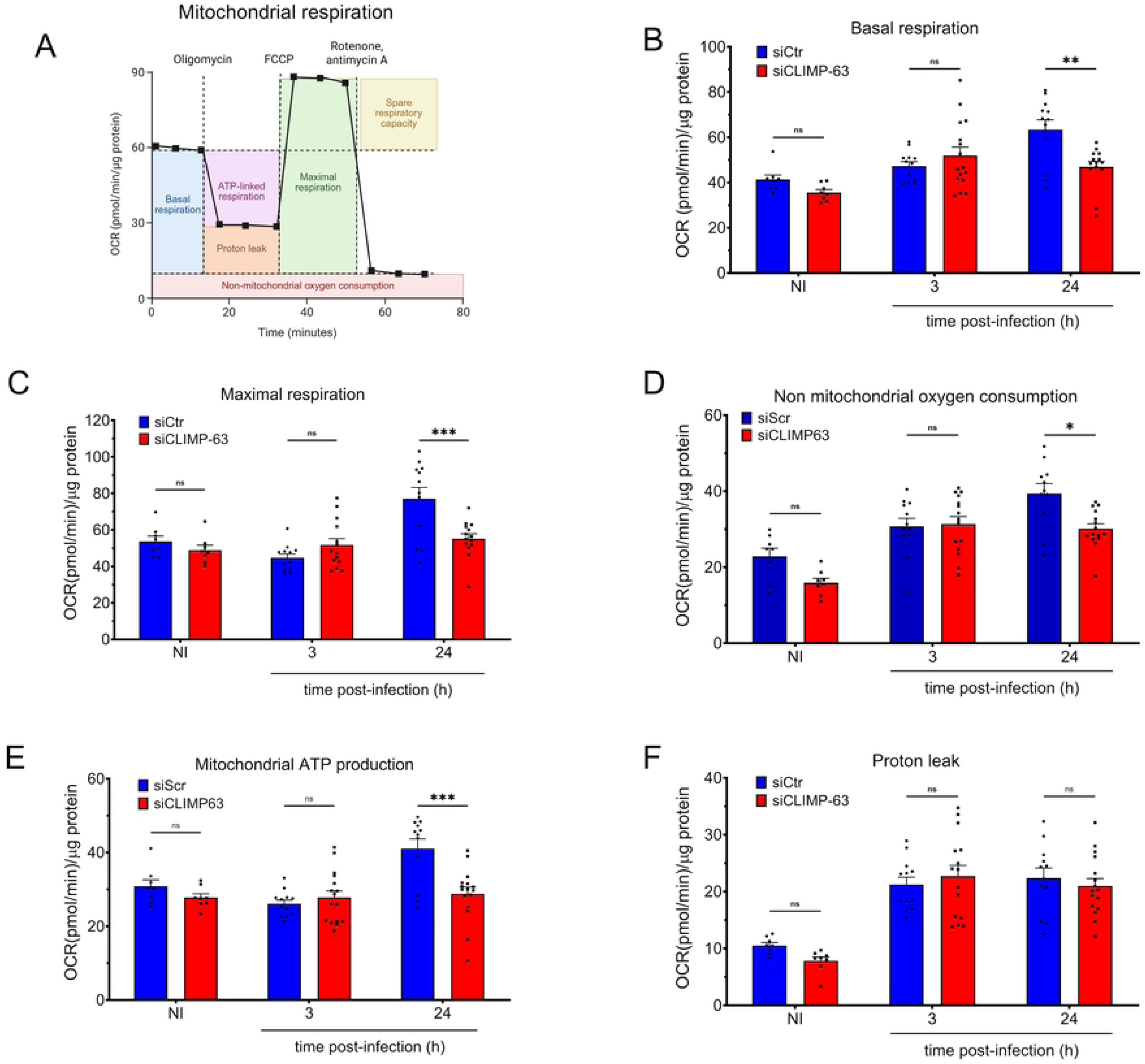
CLIMP-63 is required for *L. donovani*-induced alterations of host cell mitochondrial bioenergetics. BMM (treated with siRNA to CLIMP-63 or treated with control siRNA, Ctr) were infected or not with either *L. donovani* metacyclic promastigotes, and at the indicated time points the bioenergetic profile were determined: (A) general bioenergetic profile using the Mito Stress Test Kit, (B) basal respiration, (C) maximal respiration, (D) non-mitochondrial oxygen consumption, (E) mitochondrial ATP production and (F) proton leak. The data are presented as the means ± SEM of values of one representative experiment of three independent experiments. Statistics were calculated using one-way analyses of variance (ANOVA) with Sidak’s multiple comparison test with * P ≤ 0.05, ** P ≤ 0.01 and *** P ≤ 0.001 significance. Blots showing the efficacy the siRNA-mediated CLIMP-63 knockdowns are shown in S3 Fig.

## DISCUSSION

*Leishmania*-harboring PVs establish interactions with the mER, enabling the acquisition of host-derived components that are essential for parasite replication (1, 2, 6, 8). In this study, we investigated the contribution of the mER-shaping protein CLIMP-63 to the *Leishmania*-host cell interaction. We found that CLIMP-63 is required for the establishment and replication of *L. donovani* within macrophages and for the expansion of communal PVs harboring *L. amazonensis*. Additionally, we report that CLIMP-63 is required for *L. donovani*-induced mitochondrial biogenesis and bioenergetic metabolism remodeling, identifying this mER protein as a regulator of mitochondrial adaptation during infection. These data support a model in which *L. donovani* exploits CLIMP-63-dependent ER-mitochondria interaction to reprogram mitochondrial properties and thereby facilitate macrophage colonization.

There is a paucity of data regarding the roles of host cell ER-shaping proteins in the context of infection by intracellular pathogens. Previous studies in macrophages revealed that RTN4 is rapidly ubiquitinated and recruited to *Legionella*-containing vacuoles (LCVs), promoting bacterial replication (24, 25, 40). In contrast, CLIMP-63 associates poorly with LCVs and its depletion does not affect *Legionella* replication (25). More recently, Chopra and colleagues reported that in HeLa cells, *Salmonella* upregulates both RTN4 and CLIMP-63 transcripts; these proteins exert opposite effects on replication. RTN4 overexpression, which increases ER tubulation, led to higher *Salmonella* proliferation and more *Salmonella*-containing vacuoles (SCVs), whereas RTN4 knockdown reduces bacterial proliferation (26). Conversely, CLIMP-63 overexpression decreased the number of SCVs per cell, suggesting that increasing ER sheet abundance negatively affects *Salmonella* proliferation. They also observed that ER tubules mark the division site at the center of dividing SCVs, suggesting a role for ER tubules in vacuole fission (26). In contrast, we find that CLIMP-63, but not RTN4, is essential for *L. donovani* replication and expansion of *L. amazonensis*-harboring communal PVs, highlighting the importance of the mER for *Leishmania* to colonize macrophages. CLIMP-63 has been proposed to maintain the ER sheet structural integrity and to anchor ER membranes to the microtubule cytoskeleton (12, 13, 23, 41). The microtubule network is critical for the maturation and expansion of *L. infantum*- and *L. amazonensis*-harboring PVs, as well as for parasite differentiation and replication (42, 43). Therefore, disruption of CLIMP-63 interactions with microtubules or other ER proteins in CLIMP-63-depleted macrophages could impair LPV maturation and limit parasite replication, a possibility that warrants further investigation.

Recent studies have uncovered a novel function for CLIMP-63 in the modulation of mitochondrial biogenesis and bioenergetics. In immortalized cancer cell lines, its depletion results in a decrease in mitochondrial DNA content, compromised mitochondrial respiration, altered mitochondrial morphology and cristae structure (27, 29–31). This function is particularly relevant to *L. donovani*, as mitochondrial biogenesis and bioenergetics are critical for the ability of the parasite to colonize macrophages (32–35). Hence, pharmacological induction of mitochondrial biogenesis with AICAR significantly increases macrophage permissiveness to *L. donovani* and *L. infantum* replication, whereas co-treatment with the AMPK inhibitor compound c abrogates this effect (32, 34). Consistent with these observations, our results show that CLIMP-63 is required for *L. donovani-*induced mitochondrial DNA synthesis, ATP production, cristae remodeling, and mitochondrial respiration, providing a potential mechanistic explanation for its role in the infection process.

The *Leishmania* virulence glycolipid LPG plays a central role in the colonization of macrophages (44–49). We recently reported that LPG is essential for the ability of *L. donovani* to enhance host cell mitochondrial DNA replication and modulate mitochondrial bioenergetics (34). The identification of LPG as the factor responsible for parasite-induced CLIMP-63 redistribution provides a potential explanation for the impact of LPG on host cell mitochondria properties. Although the precise mechanism remains unclear, it is conceivable that LPG insertion into membrane bilayers and the resulting disruption of lipid rafts (36–38) contribute to CLIMP-63 redistribution following parasite internalization. Supporting this hypothesis, studies in HeLaS3 cells showed that palmitoylated CLIMP-63 maintains mitochondrial function by localizing to lipid rafts at ERMCS and binds the mitochondrial porin VDAC2 (27). CLIMP-63 binding to VDAC2 was reported to suppress interactions between VDAC2, GRP75, and IP3R, suggesting that CLIMP-63 regulates ER-mitochondria interactions by modulating ERMCS composition (50). Accordingly, dissociation of CLIMP-63 from the mitochondrial network raises the possibility that *L. donovani* alters ERMCS composition and properties, which are critical for mitochondrial dynamics and mitochondrial DNA replication (30, 31). Together, these observations support the hypothesis that *L. donovani*-induced CLIMP-63 redistribution and dissociation from mitochondria are key events underlying CLIMP-63-dependent changes in mitochondrial biogenesis and bioenergetics. Future studies will explore the mechanisms driving CLIMP-63 relocalization to LPVs, including potential effects of *Leishmania* infection on post-translational modifications that regulate CLIMP-63 trafficking, oligomerization, and function (51–53), and how these modifications contribute to macrophage colonization by *Leishmania*. Additionally, several host cell components are selectively recruited to large communal PVs and promote their expansion, including the ER-associated SNAREs Sec22b, D12, and syntaxin-5, as well as the mER membrane protein VAPA (1, 3, 8). Our finding that CLIMP-63 accumulates on communal PVs and is required for their expansion is therefore consistent with a central role of the mER in *Leishmania*-harboring communal PV development. In this regard, the differential requirement for CLIMP-63 in *L. donovani* and *L. amazonensis* may be explained by the fact that they live and replicate in two distinct types of PVs, which interact with the mER in different ways. An example of such a differential requirement for an mER protein is VAPA, which is required for the replication of *L. amazonensis* and communal PV expansion but not for *L. major* replication (8). Additional studies will be necessary to understand why CLIMP-63 is not required for *L. amazonensis* replication. Moreover, whether CLIMP-63 present on communal PVs contributes to maintaining their structure is an attractive hypothesis that warrants future investigation.

In conclusion, our work adds to the accumulating evidence that the mER plays a central role in the biology of intracellular *Leishmania* parasites. Accordingly, *Leishmania* exploits the host cell ER to create, expand, and maintain PVs, to hijack host cell resources, and to disseminate virulence factors (1, 2, 6, 8, 10). Consistent with the role of the ER in orchestrating the cellular organellar network (54–57), our findings illustrate how a pathogen manipulates the host cell ER to exploit the function and properties of other organelles.

## MATERIALS AND METHODS

### Ethic statement

Animal work was performed as stipulated by protocols 2112-01 and 2110-04, which were approved by the *Comité Institutionnel de Protection des Animaux* of the INRS-Centre Armand-Frappier Santé Biotechnologie. These protocols respect procedures on animal practice promulgated by the Canadian Council on Animal Care, described in the Guide to the Care and Use of Experimental Animals.

### Bone marrow-derived macrophages

Bone marrow-derived macrophages (BMM) were generated following established protocols (58). Marrow was harvested from the femurs and tibias of 8 to 12 week-old male and female 129/B6 mice and cultured for seven days in Dulbecco’s Modified Eagle Medium (DMEM; Thermo Fisher Scientific) supplemented with 10% heat-inactivated fetal bovine serum (FBS) (Wisent), 10 mM HEPES (pH 7.4), 100 I.U./mL penicillin, 100 μg/mL streptomycin, and 15% (v/v) L929 cell-conditioned medium as a source of macrophage colony-stimulating factor (CSF-1). Differentiation was carried out under standard conditions (37°C, 5% CO₂). To induce a quiescent state, BMM were subsequently incubated for 16 hours in CSF-1-depleted DMEM prior to experimental infection.

### Parasite cultures

Promastigotes of *L. amazonensis* LV79 (MPRO/BR/72/M1841) and *L. donovani* LV9 (MHOM/ET/67/Hu3:LV9), were maintained in *Leishmania* medium (M199 medium supplemented with 10% heat-inactivated FBS, 100 μM hypoxanthine, 10 mM HEPES, 5 μM hemin, 3 μM biopterin, 1 μM biotin, 100 U/mL penicillin, and 100 μg/mL streptomycin) at 26°C. The isogenic *L*. *donovani* LV9 Δ*lpg1* mutant (59, 60) was cultured in *Leishmania* medium supplemented with hygromycin (100 μg/mL) and its complemented counterpart *L*. *donovani* Δ*lpg1*+*LPG1* (60) was cultured in *Leishmania* medium supplemented with hygromycin (100 μg/mL) and zeocin (100 μg/mL). To preserve virulence, parasites were periodically passaged through animals and cultured for no more than six *in vitro* passages. Infective-stage metacyclic promastigotes were isolated from late stationary-phase cultures by density gradient centrifugation using Ficoll, as previously described (61).

### Infections

Metacyclic promastigotes were opsonized using C5-deficient serum obtained from DBA/2 mice, then resuspended in complete DMEM and added to BMM at a multiplicity of infection (MOI) of 7:1 for LV9 and 3:1 for LV79, following established protocols (34). To facilitate parasite-host cell contact, plates were centrifuged at 300 X *g* for 2 min using a Sorvall RT7 centrifuge. Infected cultures were subsequently incubated at 37°C, and after a 3 h period, non-internalized promastigotes were removed by three successive washes with pre-warmed Hank’s Balanced Salt Solution (HBSS).

### Antibodies

The mouse anti-CLIMP-63 monoclonal antibody (sc-393544) was from Santa Cruz Biotechnology, the rabbit anti-RTN4 polyclonal antibody (ab186735) and the rat anti-BrdU (ab6326) were from Abcam, the rabbit polyclonal antibody RTN4 (10950-1-AP) was from Proteintech, the mouse anti-phosphoglycan (Galβ1,4Manα1-PO4) CA7AE monoclonal antibody (62) was from Cedarlane, the rabbit anti-β-actin polyclonal antibody was from Cell Signalling, the rabbit anti-Tom20 polyclonal antibody (EPR15581-54) was from Abcam, and the rat anti-LAMP-1 monoclonal antibody 1D4B developed by J.T. August and purchased through the Developmental Studies Hybridoma Bank at the University of lowa and the National Institute of Child Health and Human Development. Secondary anti-mouse antibodies conjugated to Alexa Fluor 488 and to Alexa Fluor 568, secondary anti-rabbit antibodies conjugated to Alexa Fluor 647 and to Alexa Fluor 568, and secondary anti-rat antibodies conjugated to Alexa Fluor 647 were from Invitrogen-Molecular probes.

### Immunofluorescence

Cells on coverslips were fixed with 3.8% paraformaldehyde (Thermo Scientifc) for 30 min at 37°C. NH_4_Cl was used to eliminate free radicals from the cells for 10 min. Fixed cells were permeabilized for 5 min with a 0.5% Triton X-100. The samples were subsequently blocked for 1 h at 37°C with PBS containing 5% FBS prior to incubation for 2 h at 37°C with the primary antibodies diluted in PBS containing 5% FBS (1:50 for the anti-CLIMP-63, 1:200 for the anti-BrdU and the anti-LAMP1, 1:200 for the anti-Tom20 and the anti-RTN4, and 1:2000 for the anti-phosphoglycan CA7AE). Next, coverslips were incubated for 1 h at 37°C in a solution containing secondary antibodies coupled to AlexaFluor 488 (diluted 1: 500) and/or to AlexaFluor 568 (diluted 1:500) and Hoesht 33342 trihydrochloride trihydrate (1: 5000, Invitrogen, H3570) in PBS + 5% FBS). Coverslips were washed three times with 1X PBS between each step and were mounted on glass slides (Fisher) with Fluoromount-G (SouthernBiotech). Cell images were acquired on a LSM780 confocal microscope (Carl Zeiss Microimaging) using Plan Apochromat X63 oil-immersion differential interference contrast (DIC) (NA 1.64) objective, and images were acquired in sequential scanning mode. At least 30 cells per condition were processed and analyzed with ZEN 2012 software and Icy image analysis software. Control stainings confirmed the absence of cross-reactivity.

### Small interfering RNA knockdown

The siRNA transfection protocol was adapted from Dharmacon’s guidelines for cell transfection (Dharmacon). BMM were seeded in either 6-well plates, 24-well plates, or 96-well Seahorse plates from the Seahorse XF Cell Mito Stress Test kit and were reverse-transfected using Lipofectamine RNAiMAX Reagent (Invitrogen), following the manufacturer’s instructions. Transfections were carried out in a final volume of either 1000 uL, 200 μL and 30 uL of complete DMEM respectively, with a final siRNA concentration of 200 nM. BMM were either mock-transfected, transfected with a non-targeting control siRNA (Dharmacon; target sequence: UAGCGACUAAACACAUCAA), or transfected with ON-TARGETplus SMARTpool siRNAs (Dharmacon) targeting specific genes. The SMARTpool siRNA targeting *CLIMP-63* consisted of four siRNA duplexes with the following sequences: (1) CGAUGGAGUCCGACGUCUA, (2) CCGUCAAAAUCGAAACGAA, (3) GCUCAACCGAAUUAGCGAA, and (4) UCGCAGUCAGGACGUGAAA. The SMARTpool targeting *RTN4* included: (1) UGUUAUAUAUGAACGGCAU, (2) UGUAUUUGUAGGAGCGCUA, (3) UAGAUGAGCAUACUACUAA, and (4) ACAUAGAACUCCAACGUAA. Forty-eight hours post-transfection, cells were washed with complete medium and subsequently processed for downstream applications.

### BrdU labeling

BMM were seeded onto glass coverslips in 24-well plates and were infected with *L. donovani* metacyclic promastigotes and subsequently incubated at 37°C. After a 3 h period, non-internalized promastigotes were removed by three successive washes with pre-warmed Hank’s Balanced Salt Solution (HBSS). Infected cells were then incubated in complete medium supplemented with 0.1 mM 5-bromo-2’-deoxyuridine (BrdU; Sigma) for the indicated time points. Cells were then fixed with 3.7% paraformaldehyde (PFA) for 30 min at room temperature, permeabilized with 0.1% Triton X-100 for 5 min, and subsequently stained using a rat monoclonal anti-BrdU antibody (8, 39).

### Western blotting analyses

Prior to lysis, cells were washed three times with ice-cold phosphate-buffered saline (PBS) supplemented with 1 mM sodium orthovanadate, in the presence 5 mM 1,10-phenanthroline. Lysis was performed on ice using 100 μL of cold lysis buffer containing 1% triton X-100, 20 mM Hepes-KOH (pH 7.2), 150 mM KCl, 2 mM EDTA (pH 8), 1 mM dithiothreitol, 1 mM Na₃VO₄, 1 mM phenylmethylsulphonyls fluoride, 10 mM 1,10-phenanthroline and a protease/phosphatase inhibitor cocktail (Roche). Lysates were collected using a cell scraper, transferred to microcentrifuge tubes, and stored at -70°C. Prior to analysis, lysates were clarified by centrifugation for 20 min at 4°C to remove insoluble material. Protein concentrations were determined using the Pierce BCA Protein Assay Kit (Thermo Scientific), following the manufacturer’s instructions. For SDS-PAGE, 20 μg of total protein was mixed with Laemmli buffer, denatured at 100°C for 5 min, and separated by electrophoresis. Proteins were subsequently transferred to Hybond-ECL membranes, blocked for 1 h in TBS-T (0.1% Tween 20) containing 5% bovine serum albumin, and incubated overnight at 4°C with primary antibodies diluted in the same blocking buffer. Membranes were then incubated with appropriate horseradish peroxidase HRP-conjugated secondary antibodies for 1 h at room temperature. Signal detection was carried out using enhanced chemiluminescence (ECL; GE Healthcare) and visualized using the Azure Biosystem.

### Quantitative PCR analyses

To determine the mitochondrial/nuclear (mt/n) DNA ratio, total BMM DNA was extracted using a DNeasy blood and tissue kit. Quantitative PCR (qPCR) experiments were performed in three independent biological replicates and reactions were ran at least in duplicates for each sample using iTaq Universal SYBR green supermix (Bio-Rad) on a Stratagene mx3005p real-time PCR system using 10 ng DNA. The amount of mtDNA present per nuclear genome was determined using the comparative CT method (ΔΔ*CT*) (7). Relative mtDNA amounts were normalized to the hexokinase gene and expressed as the fold increase compared to non-infected controls. The DNA and RNA concentrations were determined by optical density at 260 nm (OD_260_) measurement using a NanoDrop spectrophotometer.

### Electron microscopy

For ultrastructural analysis, 5×10^6^ uninfected and *L. donovani*-infected BMM (treated with siRNA to CLIMP63 or siCtr) were fixed in 2.5% glutaraldehyde in 0.1 M cacodylate buffer (pH 7.4) overnight at 4°C. Following fixation, samples were washed three times with 0.1 M cacodylate buffer (pH 7.4) supplemented with 3% sucrose. A secondary fixation was then performed using fresh 1.3% osmium tetroxide OsO4) in collidine buffer containing 3% sucrose. Samples were dehydrated through a graded acetone series (25%, 50%, 75%, and 95%) for 30 min at each step. Following dehydration, samples were infiltrated with a 1:1 mixture of Spurr resin and acetone for 16–18 h, followed by two successive 2-hour incubations in pure Spurr resin. The samples were then processed into smaller fragments, transferred to BEEM capsules, and embedded in fresh Spurr resin. After 18 h at room temperature, the capsules were placed at 60–65°C for 30 h to allow for resin polymerization. The resulting resin blocks were cut into ultrathin sections (60–90 nm) using a Leica UC7 ultramicrotome and mounted onto Formvar- and carbon-coated 200-mesh copper grids. Sections were contrast-stained with uranyl acetate in 50% ethanol for 15 min, followed by lead citrate for 5 min. Images were acquired using a Hitachi H-7100 electron microscope equipped with an AMT camera.

### Metabolism assays

The bioenergetic profile of *L. donovani* infected BMM was analyzed using an XF-96 extracellular Flux Analyzer (Seahorse Bioscience) using the Mito Stress Test Kit. BMM were seeded at 1×10^5^ cells/well in 100 μl of DMEM in XF-96 cell culture plates and reverse-transfected following the Small interfering RNA knockdown mentioned above. After, BMM were infected with *L. donovani* metacyclic promastigotes at the indicated time post-infection. One hour before the readouts, the cells were washed and the medium was changed to XF medium (buffered DMEM supplemented with 4.5 g/L of glucose, 2% of FBS, 2 mM L-glutamine, 100 U/ml penicillin and 100 mg/ml streptomycin). The real-time measurement of the bioenergetic profile was obtained under basal conditions and in response to oligomycin (1μM), fluoro-carbonyl cyanide phenylhydrazone (FCCP 2μM), Rotenone (1μM) and Antimycin A (1μM). The non-mitochondrial respiration was obtained by subtracting the Rotenone/Antimycin A values. The procedure used in the experiments was established according to Seahorse manufacturer instructions.

### Statistics and reproducibility

Graphs and statistical analyses were performed using GraphPad Prism version 11. Each experiment was independently conducted three times. Specific statistical tests, exact sample sizes (n), and the nature of error bars are detailed in the respective figure legends. Statistical significance is indicated as follows: * P<0.05, ** P<0.01, *** P<0.001 and **** P<0.0001 according to one-way analysis of variance (ANOVA) with a Sidak’s or Tukey’s multiple comparison test. All immunoblots and microscopy images presented are representative of at least three independent experiments.

## Acknowledgments

We thank Mr. Jessy Tremblay (Confocal microscopy and flow cytometry Facility, Centre Armand-Frappier Santé Biotechnologie) for expert assistance in immunofluorescence confocal microscopy and Mr. Arnaldo Nakamura (Electron microscopy Platform, Centre Armand-Frappier Santé Biotechnologie) for expert assistance in electron microscopy.

## Author contributions

Conceptualization: Charles-Antoine Boyer, Hamlet Acevedo Ospina, Albert Descoteaux.

Data curation: Charles-Antoine Boyer, Hamlet Acevedo Ospina,, Albert Descoteaux.

Formal analysis: Charles-Antoine Boyer, Hamlet Acevedo Ospina,, Albert Descoteaux.

Funding acquisition: Albert Descoteaux.

Investigation: Charles-Antoine Boyer, Hamlet Acevedo Ospina,.

Methodology: Charles-Antoine Boyer, Hamlet Acevedo Ospina,, Albert Descoteaux.

Project administration: Albert Descoteaux.

Resources: Albert Descoteaux.

Supervision: Albert Descoteaux.

Visualization: Charles-Antoine Boyer, Hamlet Acevedo Ospina,, Albert Descoteaux.

Writing – original draft: Charles-Antoine Boyer, Albert Descoteaux.

Writing – review & editing: Albert Descoteaux.

## Supporting information

**S1 Fig. Distribution of RTN4 in BMM infected with *L. donovani* and *L. amazonensis***. BMM were infected or not (NI) with either (A) *L. donovani* or (B) *L. amazonensis* metacyclic promastigotes and at the indicated times points, the distribution of RTN4 (red) was assessed by confocal immunofluorescence microscopy. DNA is in blue. White arrowheads denote internalized parasites.

**S2 Fig. CLIMP-63 is required for the replication of *L. donovani* in BMM**. (A) Labeling of *L. donovani* amastigotes with BrdU in BMM (treated with control Ctr siRNA or siRNA to CLIMP-63) infected for 48 h and 72 h. (B) Percent of BrdU+ parasites in BMM (treated with control Ctr siRNA or siRNA to CLIMP-63) at 48 h and 72 h post-phagocytosis.

**S3 Fig. Efficacy of siRNA-mediated knockdown of CLIMP-63 in BMM.** Representative Western blots of CLIMP-63 levels in BMM treated with control siRNA and in BMM treated with siRNA to CLIMP-63. Levels of β-actin were used as controls. Blots for the levels of CLIMP-63 for the results shown in Figure 3 (A-C), Figure 4 (B-E), and Figure 5.

**S4 Fig. Efficacy of siRNA-mediated knockdown of RTN4 in BMM.** Representative Western blots of RTN4 levels in BMM treated with control siRNA and in BMM treated with siRNA to RTN4. Levels of β-actin were used as controls. Blots for the levels of RTN4 for the results shown in Figure 3 (D-F).

**S1 Data. Source data for Figs 3, 4, 5, S2B**

## Data availability statement

All relevant data are within the manuscript and its Supporting Information files.

